# Spurious correlation inflates performance in single-cell perturbation prediction

**DOI:** 10.64898/2026.05.07.723486

**Authors:** Phillip B. Nicol, Shriya Shivakumar, Rafael A. Irizarry

## Abstract

The increasing number of computational methods designed to predict the effects of genetic perturbations on cellular gene expression profiles has led to a need for rigorous evaluation metrics. Recent benchmarking studies rely on correlation or cosine similarity of differential expression relative to a shared population of control cells. We show that these metrics are systematically inflated by statistical bias induced by reusing the same control population to define both quantities being compared. As a result, even non-informative methods can appear to perform well, particularly in datasets with limited numbers of control cells. Reanalysis of published datasets using a simple control-splitting procedure that removes this bias leads to a substantial reduction in performance previously attributed to biological signal.

---

There is increasing interest in developing computational tools to predict the effects of genetic perturbations, such as gene knockouts or CRISPR edits, on cellular gene expression profiles. The general approach is to train models on gene expression data from a collection of perturbed cells and then predict the transcriptional response to unseen perturbations, given only the identity of the perturbation. The proposed approaches differ substantially in methodology, ranging from deep learning models, such as scGPT (Cui et al., 2024), GEARS (Roohani et al., 2024), and STATE (Adduri et al., 2025), among others (Wei et al., 2026), to much simpler statistical strategies (Ahlmann-Eltze et al., 2025). In particular, recent studies have shown that extremely simple baselines, such as predicting any perturbation by the mean expression profile across the training data, can perform competitively in this task (Viñas Torné et al., 2025).

Given the number and range of proposed methods, often reported to achieve strong performance, it is essential to use rigorous evaluation metrics to assess predictive performance. Commonly used reference-sensitive metrics include the cosine similarity or Pearson correlation of differential expression vectors relative to a population of control cells (denoted with Pearson_Δ_), as these prioritize the directional accuracy of the prediction (Viñas Torné et al., 2025; Wei et al., 2026). However, the statistical properties and limitations of these metrics have only begun to be explored (Liu et al., 2025; Vollenweider and Buehlmann, 2026; Mejia et al., 2025). Here, we show that the standard practice of using the same control cells to estimate both the true and predicted expression vectors induces *spurious correlation*, inflating apparent model performance. We show that this bias can be substantial, particularly in datasets with a small number of control cells. Notably, the risk of spurious correlation arising from shared random variables has been understood for over a century (Pearson, 1897), yet its implications for modern evaluation practices in this setting appear to have been overlooked.

Specifically, to demonstrate the practical consequences of this bias, we construct a non-informative prediction method that, by design, ignores both the training data and the identity of the perturbation. Despite containing no perturbation-specific information, this method achieves performance comparable to leading models under current evaluation metrics. The apparent competitive performance of this method vanishes when two independent groups of control cells are used to form the reference. A reanalysis of several published datasets shows that a substantial fraction of reported performance can be attributed to this bias, and that qualitatively different conclusions are reached when using a corrected evaluation.

To explain in detail the source of the bias, consider a single perturbation that results in a mean (normalized) expression *µ*_1_ ∈ ℝ^*d*^, with *d* the number of genes being measured. Let *µ*_0_ ∈ ℝ^*d*^ denote the mean expression of a control cell. A perturbation prediction algorithm produces a predicted expression profile 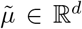 for *µ*_1_, and we condition on this prediction in the analysis that follows. The expression changes relative to the control is defined as Δ_1_ = *µ*_1_ − *µ*_0_ and 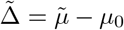. The Pearson_Δ_ (denoted here *ρ*_Δ_) metric is defined as the Pearson correlation of the entries of Δ_1_ and 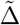. Assuming the entries are mean centered, this can be written as

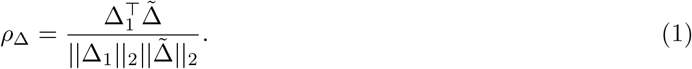

Because *µ*_0_ and *µ*_1_ are unknown parameters, it is not possible to compute (1) exactly. Instead, an estimator 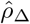 is computed by plugging in the sample means (centroids):

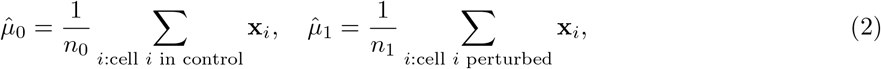

where **x**_*i*_ ∈ ℝ^*d*^ denotes the observed *d*-dimensional vector of normalized expression values for cell *i*, and *n*_1_ and *n*_0_ represent the perturbed and control sample sizes, respectively. However, although this plug-in approach appears natural, the resulting estimator 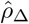 is biased upward relative to the true value *ρ*_Δ_ because 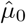 is used in both terms of the numerator, inducing dependence through shared randomness. Ignoring this effect leads to systematically overoptimistic conclusions about model performance.

This type of induced correlation is a direct instance of the spurious correlation effect first described by Karl Pearson:

> “If *u* = *f*_1_(*x, y*) and *v* = *f*_2_(*z, y*) be two functions of the three variables *x, y, z*, and these variables be selected at random so that there exists no correlation between *x, y, y, z*, or *z, x*, there will still be found to exist correlation between *u* and *v*. Thus a real danger arises when a statistical biologist attributes the correlation between two functions like *u* and *v* to *organic relationship*.”
>
> — Pearson (1897)

To demonstrate, we simulated 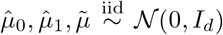 and observed 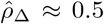, despite negligible correlation in the entries of 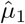 and 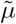 (**Fig 1a**). Although the upward bias is expected to decay as *n*_0_ increases, we show in Proposition 1 that it is substantial in the high-dimensional settings typical of single-cell data, where expression profiles span hundreds to thousands of genes. As a result, the observed metric 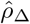 will appear substantially closer to 1 than *ρ*_Δ_. Moreover, the bias is typically larger when *µ*_0_ and 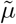 are close, meaning that 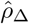 can spuriously favor methods that produce predictions close to the control cells. A simple approach to mitigate this bias is to compute the Pearson correlation of 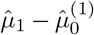 and 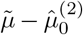, where 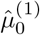 and 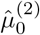 are formed from two randomly chosen (independent) groups of control cells. We refer to this estimator as 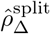 and 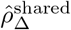 as the (standard) estimator described above.

**Figure 1:**
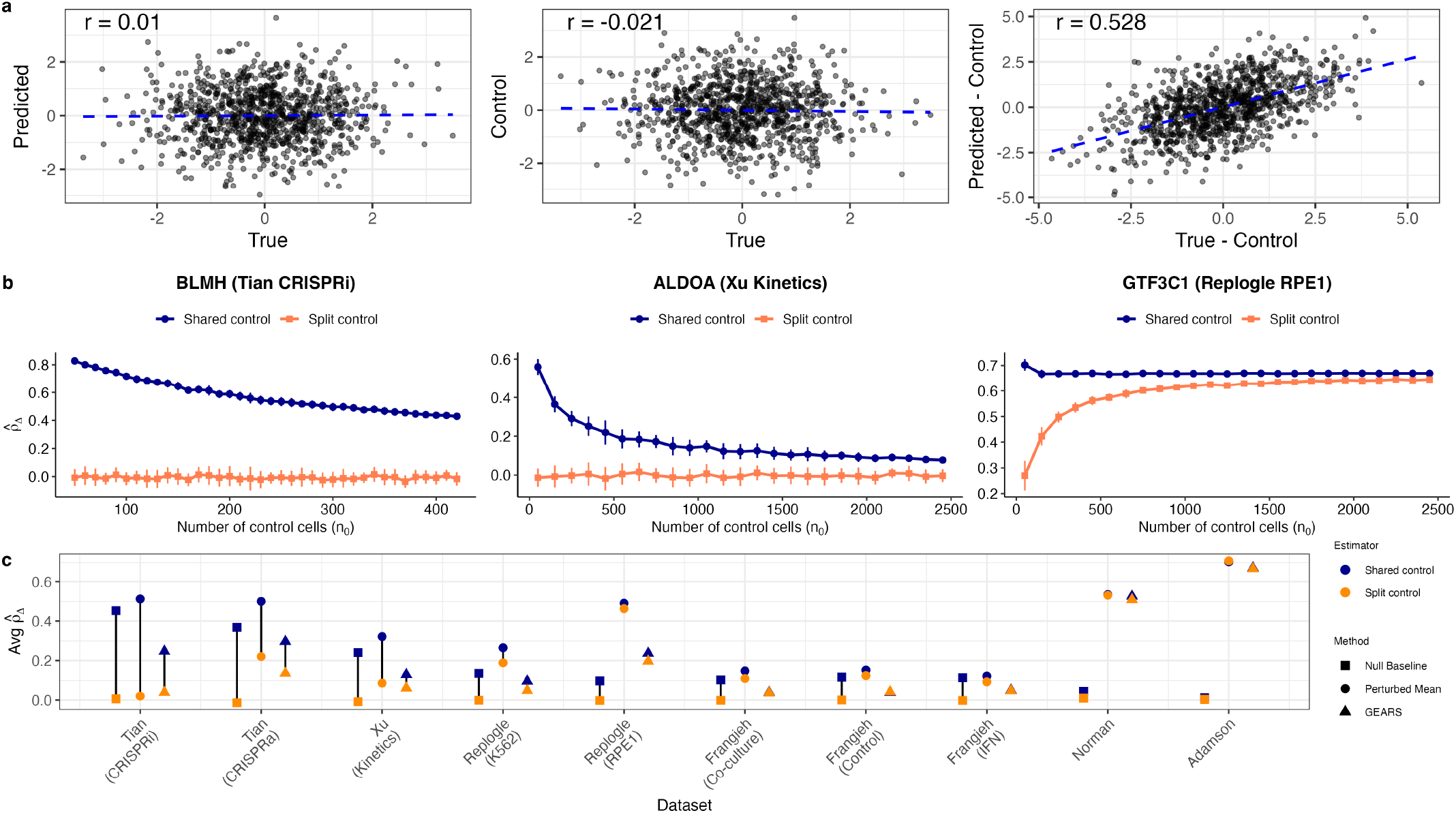
a. We simulated the true, predicted, and control expression for each of *d* = 1000 genes from a *N* (0, 1) distribution. Although there is negligible correlation between the true and predicted expression (left panel), subtraction of the control expression induces strong positive correlation (right panel). **b**. We selected three perturbations from three different datasets and compared 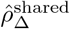 and 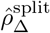 as the number of control cells *n*_0_ used to form the reference varies. **c**. The average 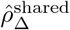 and 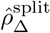 across all perturbations in 10 datasets. Three methods were used to produce 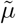 (1) the perturbed mean method, (2) a null baseline that sets 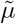 to be the mean of held-out control cells, (3) GEARS.

To quantify the impact of this bias, we evaluated three representative perturbations across three datasets, comparing the standard evaluation metric 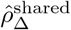 with 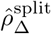. As a representative example, we considered the simple baseline predictor identified as the top-performing method by Viñas Torné et al. (2025), setting 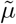 to the mean expression of perturbed cells in the training data, and varied the number of control cells *n*_0_ used to define the reference. Under the standard approach, 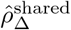 reached values as high as 0.8 for the Tian et al. (2019) and Xu et al. (2024) datasets (**Fig 1b**), suggesting strong predictive performance. In contrast, the corrected estimator 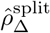 remained near zero across all values of *n*_0_, indicating no detectable predictive signal. As *n*_0_ increased, the apparent performance reported by 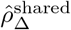 collapsed towards zero, consistent with substantial upward bias at smaller sample sizes.

This discrepancy is not subtle: apparent high correlations can be obtained even in the complete absence of perturbation signal. In particular, correlation estimates exceeding 0.75 arise when comparing two groups of cells drawn entirely from the control population (**Fig S1**). Consistent with this, principal component projections show no discernible separation between control and perturbed cells in the Tian et al. (2019) and Xu et al. (2024) datasets (**Fig S2**), indicating that the apparent predictive accuracy under 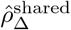 is not supported by underlying biological signal.

To demonstrate the practical consequences, we constructed a null baseline by splitting the control cells into two equal-sized groups, using the first to define 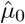 and the second as the prediction 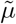. Despite not using any data from the perturbed cells, this method achieves 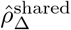 values that match or exceed those of GEARS (Roohani et al., 2024) on several datasets. When evaluated with 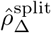, however, its performance collapses to zero, as expected. Extending this analysis across all perturbations in the ten datasets studied by (Viñas Torné et al., 2025), we find that the majority of reported performance in the Tian et al. (2019) and Xu et al. (2024) datasets disappears under the corrected evaluation metric (**Fig 1c**). These results show that 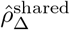 can both inflate absolute performance and distort the relative ranking of prediction algorithms.

Although the control-splitting estimator removes the upward bias, it is typically conservative, with estimates attenuated toward zero (Proposition 2). This is evident for the perturbation from Replogle et al. (2022) (RPE1) in Figure 1b, which appears to be genuinely well predicted by 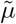. However, we view this conservative approach as preferable, as it avoids spurious overestimation of performance and can still rank methods correctly despite underestimation of true correlation. In practice, the split estimator can be further stabilized by averaging over multiple random splits (Methods).

A commonly used variant of *ρ*_Δ_ instead computes the correlation restricted to the top *k* differentially expressed genes (DEGs) between the perturbed and control cells. This metric aims to prioritize recovery of the most important changes induced by the perturbation. A common choice is *k* = 20, in which case the metric is known as Pearson Δ_20_ (Viñas Torné et al., 2025). Despite the restriction to *k* ≪ *d* genes, this metric can still suffer from substantial upward bias if the same set of cells is used to select the *k* DEGs and to compute the correlation. Returning to the simulated null example from Figure 1a, we recomputed the correlation on the subset of genes with the 20 largest effect sizes and observed *r* = 0.87 (**Fig 2**a), demonstrating that the selection of top DEGs can amplify the spurious correlation. To mitigate this bias, we first estimate the top *k* DEGs using an independent set of cells, and then to use the remaining cells to compute 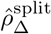 restricted to the top *k* indices (see Methods for additional details). On the real datasets, we again tend to observe significant drops in the estimated correlation when using the splitting estimator (**Fig 2**b). As before, we also observe changes in the relative ranking of methods; for example, the standard shared-data estimator ranks the perturbed mean above GEARS on the Xu et al. (2024) dataset, whereas the ordering reverses when the split-based estimator is used.

**Figure 2:**
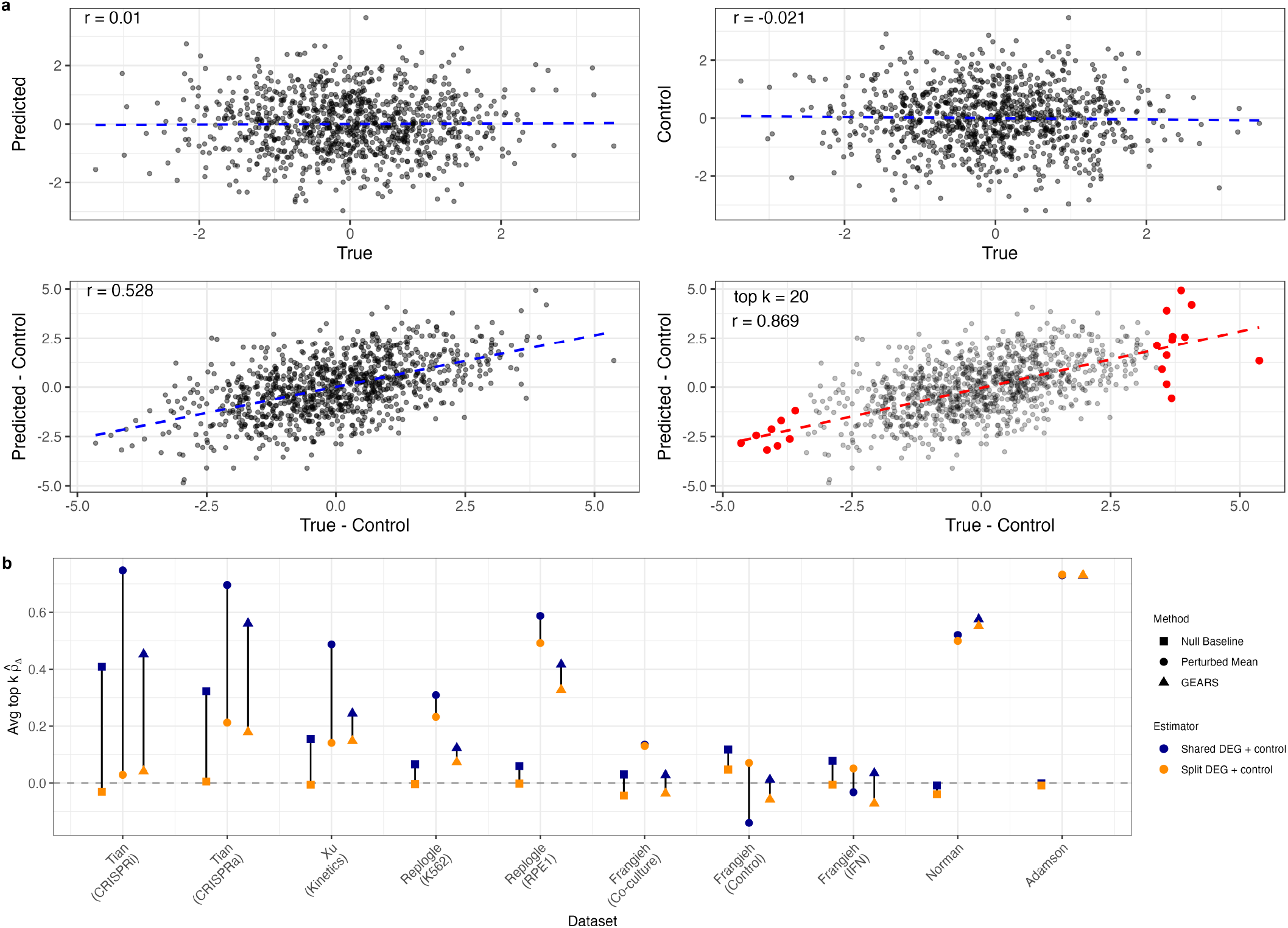
a. We extended the simulation in Figure 1a to compute the correlation of the top *k* differentially expressed genes (DEGs). Here, the top *k* DEGs are selected as the *k* largest values of 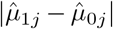, 1 ≤ *j* ≤ *d*. **b**. We repeated the analysis of Figure 1c for the top *k* Pearson Δ, comparing the average correlations when using the shared or splitting-based estimator.

The central contribution of our note is to recognize that perturbation prediction metrics are implicitly point estimates of population parameters. By treating the performance metric as an estimated parameter, we gain a framework for identifying and correcting the statistical biases influencing performance. Similar approaches can be useful for designing new robust and principled metrics to compare the ever-increasing number of prediction methods.

## Methods

### Theoretical analysis

Here, we compute the asymptotic behavior of the shared and split estimators of Pearson_Δ_. We will make the following proportional asymptotic assumption:

#### Assumption 1 (Proportional asymptotics)

*n*_0_, *n*_1_, *d* → ∞ *in such a way that d/n*_0_ → *c*_0_ ∈ [0, ∞) *and d/n*_1_ → *c*_1_ ∈ [0, ∞).

This asymptotic framework is well-suited for describing single-cell genomic data; for instance in the datasets we considered *d* ≈ 5000 and *n*_0_ ranged from approximately 500 (*c*_0_ = 10) to 25000 (*c*_0_ = 0.2). We assume that noise in control cells is uncorrelated across genes and has equal variance:

#### Assumption 2 (Noise)

*The control cells* **x**_1_, …, 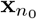 *are i*.*i*.*d. with mean µ*_0_, *covariance σ*^2^*I*_*d*_, *independent coordinates* **x**_*ij*_, *and uniformly bounded central fourth moments:*

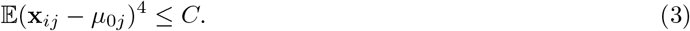

*The perturbed cells have mean µ*_1_ *and the same second and fourth moment assumptions*.

For simplicity, we will condition on 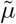 and therefore treat it as a deterministic vector (i.e., the limits will be conditional on 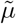).

#### Assumption 3 (Signal strength)

*For each n*_0_, *µ*_0_, *µ*_1_, 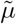 *are deterministic vectors such that*

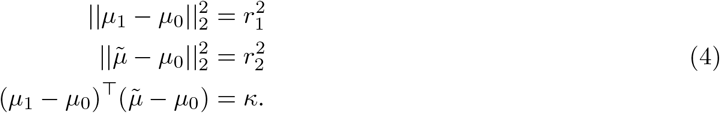

Under the above assumption, the true cosine similarity (Pearson_Δ_ for centered vectors) may be defined as

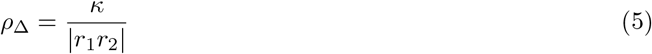

The two considered estimators are

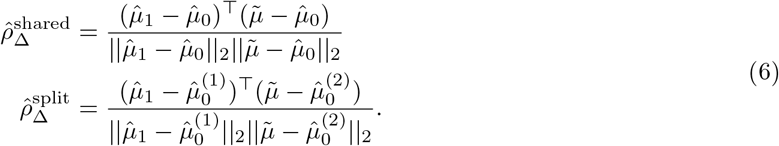

##### Proposition 1

*Under assumptions 1, 2, 3, we have*

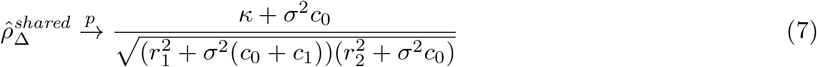

If we assume 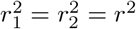, *c*_1_ = 0 then the above limit may be written as

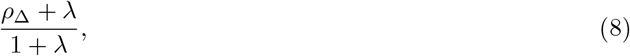

where *λ* = *σ*^2^*c*_0_*/r*^2^. In this case, the limiting bias is strictly positive (provided *ρ*_Δ_ *<* 1) and decreases as *n*_0_ increases. From this, we can also see that the bias is larger when 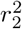 is close to 0. This provides intuition for why the hold-out mean in Figure 1b appears unreasonably strong. Simulations suggest the theoretical predictions are accurate when *d* ≈ 5000 (**Fig S3**). On the contrary, 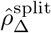 removes the numerator bias:

##### Proposition 2

*Under assumptions 1, 2, 3, we have*

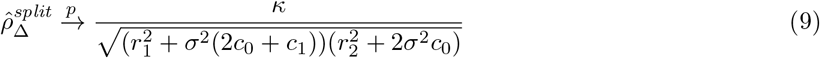

The previous proposition makes the difference between 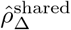 and 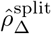 immediately clear. Instead of a persistent upward bias, 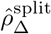 is instead increasingly biased towards the null (i.e., it is biased downwards when the true *ρ*_Δ_ *>* 0). Most importantly, 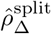 is consistent under the null whereas 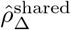 is not.

### Practical estimator

In practice, 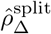 should be computed as the average of multiple random splits. Let *B* be the number of random splits (we use 100), and define 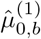 and 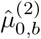 to be the control means computed from the two random groups on the *b*-th split. Then

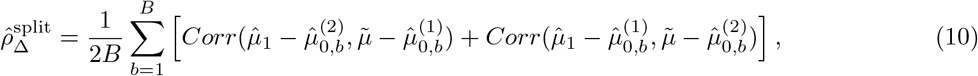

where *Corr*(·,·) denotes the Pearson correlation coefficient of the two vectors (cosine similarity after centering). This form of the estimator was used to measure the performance of the perturbed mean and GEARS in Figure 1c.

### Correlation restricted to top differentially expressed genes

Here we provide the step-by-step details of the splitting approach to compute the top-*k* Pearson Δ:

1. Within each perturbation (including controls), equally divide the cells into two groups. Across all perturbations, this produces two subsets of cells *A* and *B*.
2. On subset *A*, select the top *k* DEGs (for each perturbation). Let *Ŝ*_*k,p*_ denote the selected genes for perturbation *p*. Although any selection procedure could be used in this step, in Figure 2b we use the approach in the GEARS python package to obtain the list of DEGs.
3. On subset *B*, apply 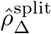 (Equation (10)) restricted to entries *Ŝ*_*k,p*_.

We note that there is some ambiguity in what the corresponding population analogue of the top *k* Pearson Δ. For example, one could define *S*_*k*_ to be the subset corresponding to the *k* largest values of |*µ*_1*j*_ − *µ*_0*j*_ |, 1 ≤ *j* ≤ *d*, but there are other possibilities as well. We also believe that considerable improvements could also be made to our proposed approach; for instance, by varying the sizes of subsets *A* and *B*.

## Data and code availability

We performed our analysis within the Systema framework (Viñas Torné et al., 2025), using their code (Viñas and Jiang, 2025) to process and evaluate the different datasets. The considered datasets were originally generated by Tian et al. (2019), Xu et al. (2024), Replogle et al. (2022), Frangieh et al. (2021), Norman et al. (2019), Adamson et al. (2016). For Replogle et al. (2022) and Norman et al. (2019), we downloaded the data from scPerturb (Peidli et al., 2024) and observed that downstream results varied only slightly from the results of (Viñas Torné et al., 2025).

Our fork of their repository is available at: https://github.com/phillipnicol/systema.

## Acknowledgements

The authors thank Isabella Grabski, Tavor Baharav, Eric Weine, and Phuc Vu for helpful discussion. PBN was supported by the National Institutes of Health grant T32CA009337.

## Supplementary Material

### S1 Proofs

The following is a direct application of the Chebyshev inequality:

#### Lemma 4

*Under assumptions 1, 2, 3*,

1. *If a*^(*d*)^ *are a sequence of deterministic vectors with* 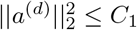 *for all d, then*

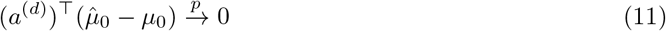

2. 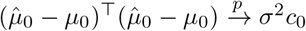

3. 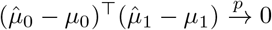

#### Proof of Proposition 1

Using Lemma 4, the numerator converges to

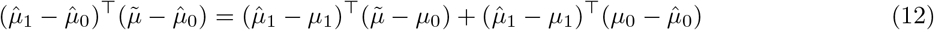

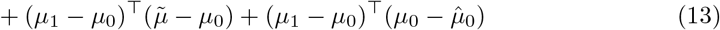

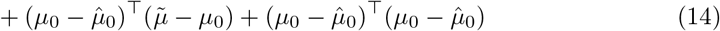

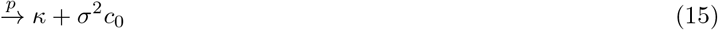

On the denominator, use a similar idea:

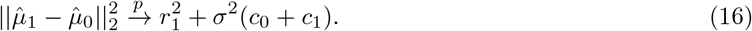

Conclude by applying the continuous mapping theorem.

#### Proof of Proposition 2

Because the controls are split into two equal sized groups (assume *n*_0_ even), the denominator converges to the same value as in Proposition 1 with 2*c*_0_ replacing *c*_0_.

Let 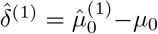 and 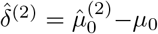. The result for the numerator follows provided that 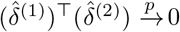. First, by independence

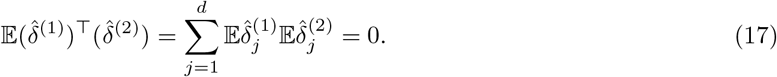

For the variance, see that

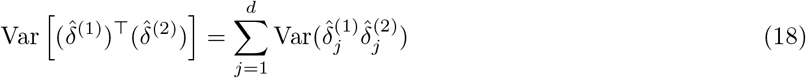

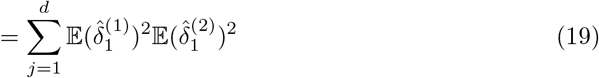

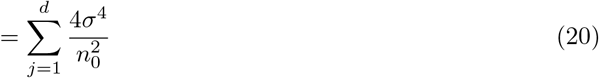

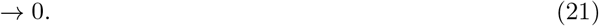

### S2 Supplementary Figures

**Figure S1:**
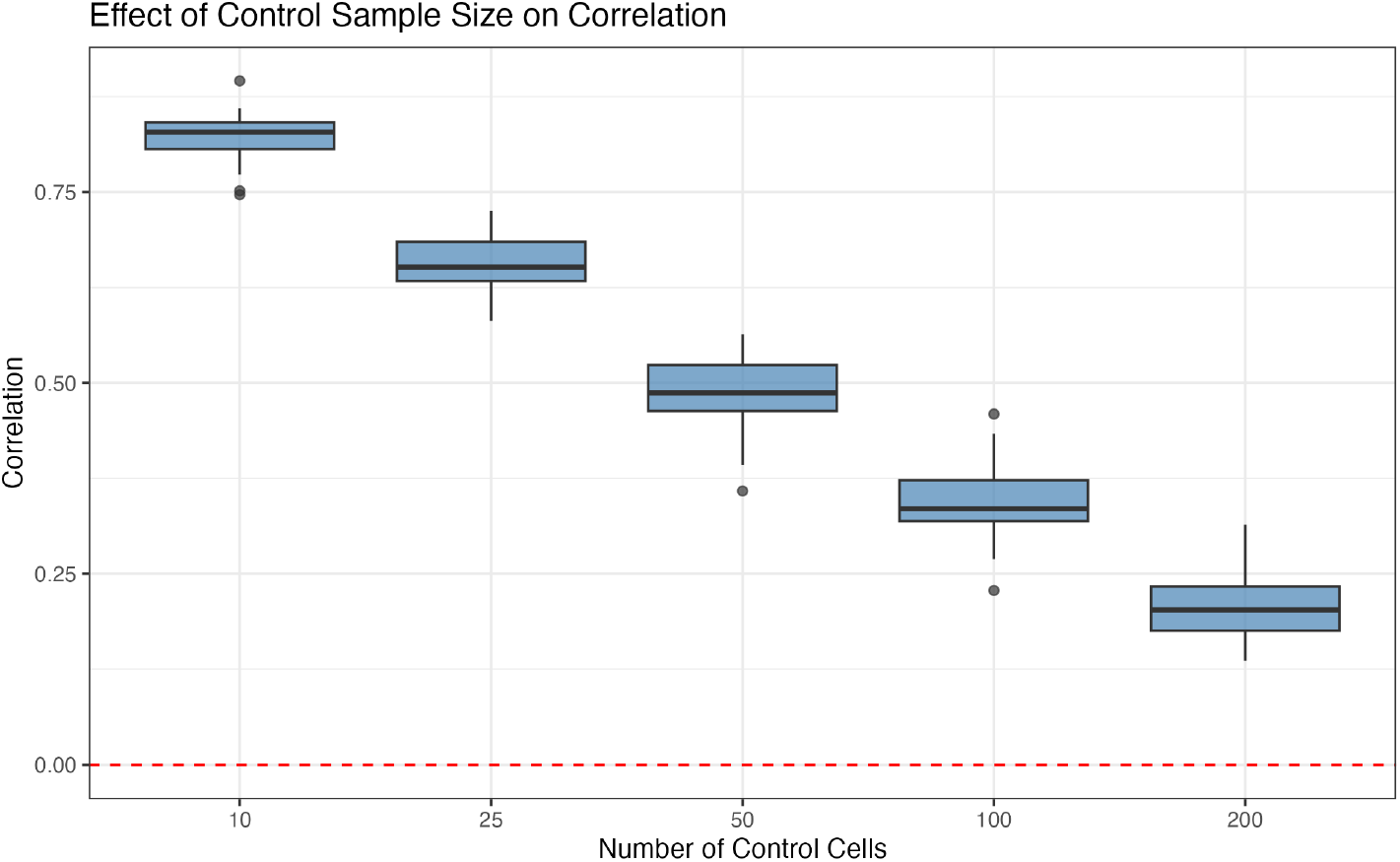
Using only the control cells from the Tian et al. (2019) CRISPRi dataset, we sampled *n*_0_ cells to be the “control”, *n*_1_ = 50 cells to be the perturbed, and another *n*_1_ cells to construct the estimate of 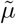.

**Figure S2:**
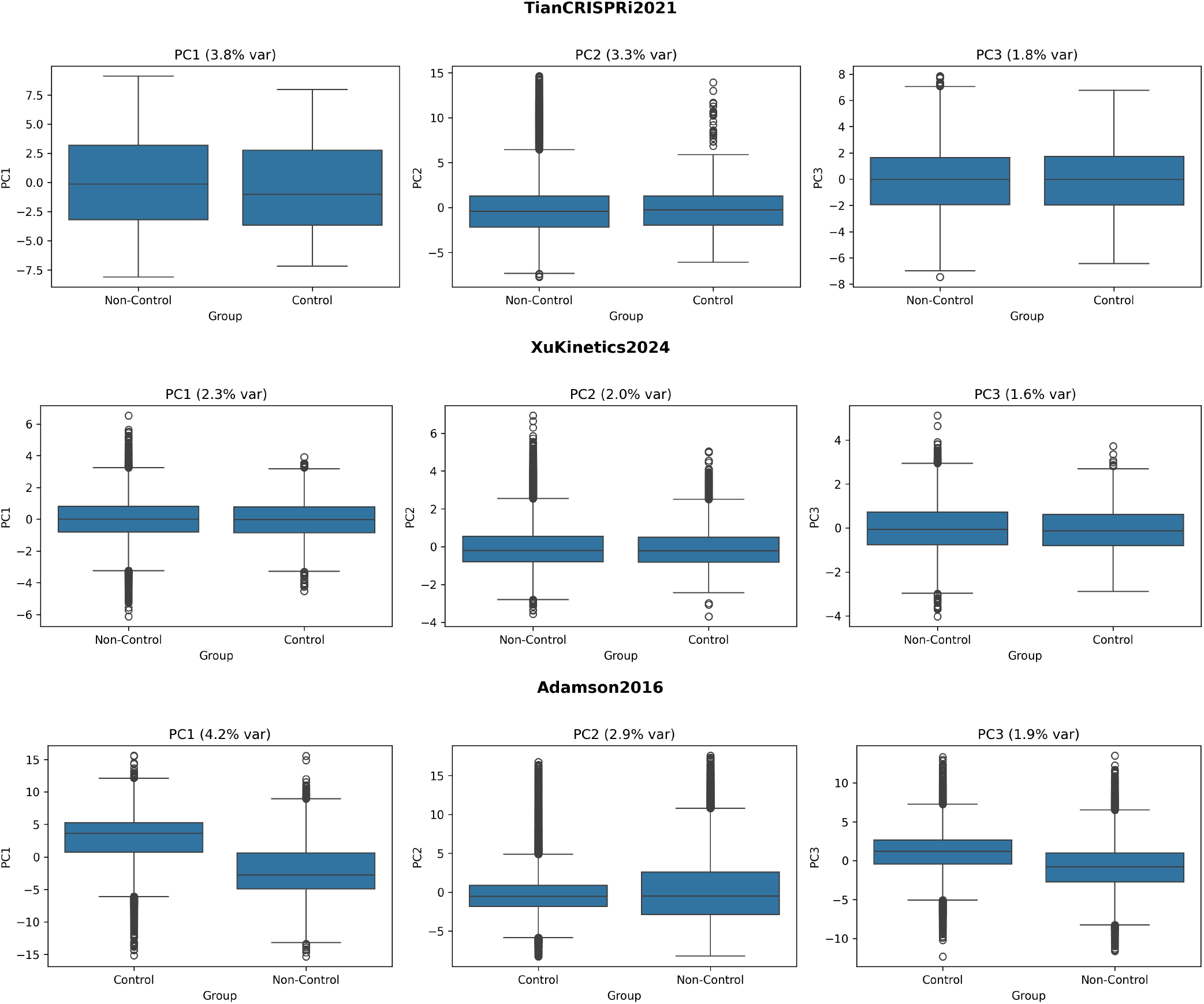
Boxplots of the first three principal components stratified by control status (for three different datasets).

**Figure S3:**
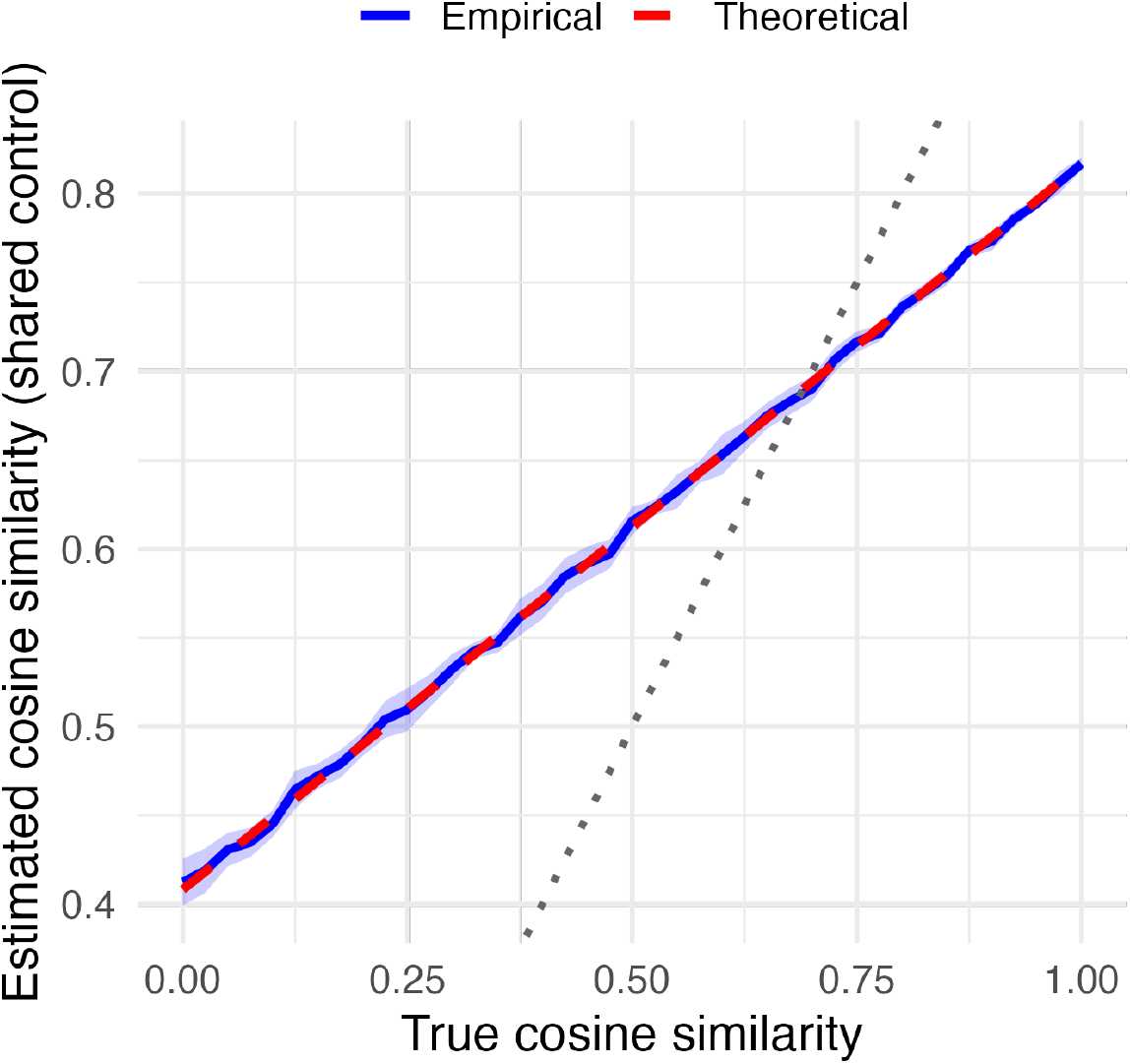
Simulated cosine similarities vs the theoretical value in Proposition 1. We drew expression values from a multivariate normal distribution with *σ*^2^ = 1, *n*_1_ = *n*_0_ = *d* = 5000, *µ*_1_ = *e*_1_, *µ*_0_ = 0, 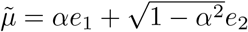 (with variable *α*).

## Notes

### Competing Interest Statement

The authors have declared no competing interest.

### Summary of Updates

Adding analysis of Pearson Delta on top k genes.

